# Amino acid repeat mosaics shape protein functional landscape

**DOI:** 10.64898/2026.06.09.730932

**Authors:** Anjali Kumari Singh, Anurag Babu, Adhithya Subramanian Gopalakrishnan, Nagashree Rachote, Vaidehi Sharma, Sourav Ganguli, Keertana Sai Kappagantula, Harikrishnan Ramadasan, Palak Shukla, Arunkumar Dhayalan, Pavithra L Chavali, Sreenivas Chavali

**Author notes:** These authors contributed equally: Anurag Babu, Adhithya Subramanian Gopalakrishnan.

## Abstract

How combinations of functional units such as repeats and motifs within disordered regions influence protein functions remains poorly understood. Here, we investigate how co-occurring amino acid homorepeats of different types within the same protein (HR-mosaics) shape molecular outcomes. Using a novel evolutionary co-occurrence metric, human HR-mosaics were classified as segregated (each HR independently influences distinct outcomes), concerted-disjunct (both HRs jointly shape protein functionality without modulating each other), or concerted-conjunct (both HRs affect functionality through mutual influence). Molecular studies of naturally occurring polyGly-polyPro mosaic in DDX17 and chimeric mosaics of polyAla and polyHis demonstrate that HR-mosaics influence protein abundance, localisation, mobility and interaction landscape. Segregated HR-mosaics expand functional space, concerted-disjunct additionally confer biological context, and concerted-conjunct mosaics further enable rheostatic regulation between HRs. These design principles underscore amino acid type, length and relative positioning of HRs as central architects of HR-mosaic functionality, with implications in protein design and engineering and understanding repeat-associated pathologies.

## Introduction

Functional units, such as structural domains, contribute to the molecular function(s) of proteins either as modular units or in combinations. Importantly, multi-domain architectures provide an ensemble of molecular functions to a protein^1–12^. Novel combinations of such functional units arise during evolution, as a consequence of gene duplication, sequence divergence and/or gene fusion^2,5^. Such coalition and shuffling of functional units in proteins is associated with evolution of protein functionality^2–4^. This enables greater proteome complexity and exploration of a broader functional space. However, there is a limited understanding of how combination-based assortment of functional units emerge and contribute to functional outcomes in intrinsically disordered regions (IDRs) of proteins.

Growing evidences indicate that homorepeats (HRs), which are stretches of identical amino acids (e.g. polyglutamine) serve as important functional units within IDRs. HRs contribute to a wide range of molecular functions, including modulation of subcellular localisation, biomolecular interactability and transcriptional regulation^13–17^. Molecular studies have shown functional associations between two different amino acid HRs within the same protein. For instance (i) the presence of polyPro HR at the C-terminus of polyGln in a chimeric polypeptide reduces protein aggregation that could be driven by improper conformation of polyGln, and enables stable physiological interactions with coiled-coil partners^18,19^, and (ii) the length ratio of polyGln to polyAla HRs in RUNX2 is linked to the cranial morphology of canines^20^. Do combinatorial HRs emerge from specific configurations of individual amino acid type HRs? Preliminary studies show that certain HR combinations are conserved in developmental proteins, such as the HOX, FOX and SOX proteins. For instance, polyAla and polyGly co-occurrences, and polyArg and polyPro co-occurrences are conserved across metazoan proteins involved in skeletal development and heart development, respectively^21^. These studies suggest that HR combinations could be associated with protein functional outcomes. These observations raise fundamental questions in protein design: How do combinations of repeats affect protein functions, and what are the distinct modes by which HRs operate in combinations? Through extensive analysis of the human proteome in comparison with other eukaryote proteomes, and experimental investigations of human and chimeric proteins, we present a comprehensive framework that elucidates the roles and possible modes of actions of HR combinations in affecting protein function(s).

## Results

We first identified human proteins with amino acid HRs (HRPs), defined as a protein with at least one continuous stretch of five or more identical amino acids. On the basis of the number and type of amino acid HRs present, we then classified human HRPs into (i) single HR containing HRPs (singleton HRPs), (ii) multiple HR containing proteins with all HRs of the identical amino acid type (similar HRPs) and (iii) multiple HR containing proteins with at least two different amino acid HRs (dissimilar HRPs) [**Fig. 1A; Supplementary Fig. 1A**]. Interestingly, human proteome is enriched for having multi-HR containing proteins (similar and dissimilar HRPs) [**Supplementary Fig. 1B**]. Furthermore, even within multi-HRPs, dissimilar HRPs are more enriched compared to singleton HRPs. While, ∼30% of human HRPs contain more than one HR, 72% of these multiple HR containing proteins are constituted by dissimilar HRPs [**Supplementary Fig. 1C**].

**Fig. 1.**
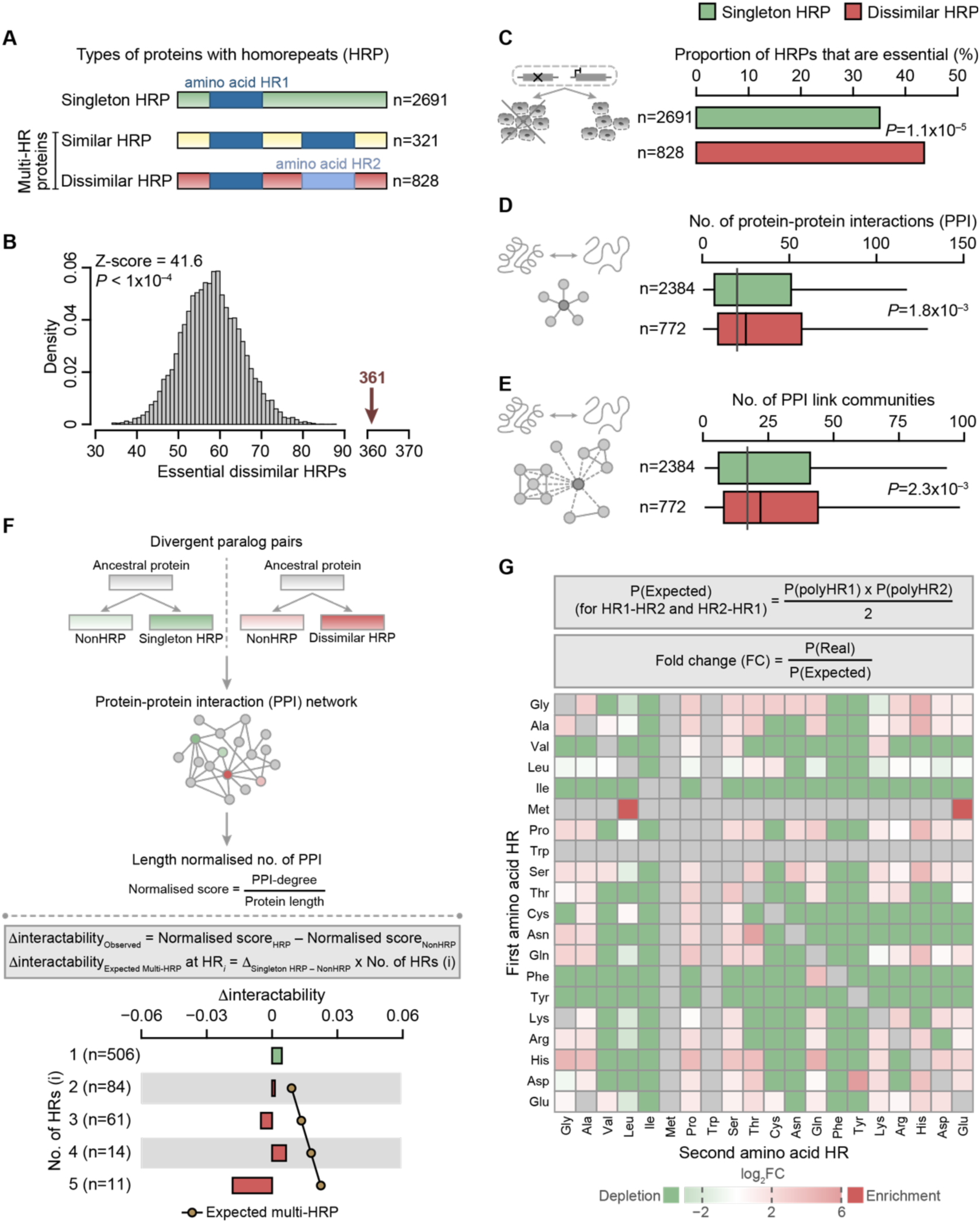
Proteins harboring dissimilar HRs are physiologically important, exhibit a disproportionate shift in interaction capacity and a non-random organizational pattern of the constituent HRs. (A) Classification of the human proteins with homorepeats (HRP) based on the number and type of HRs in each protein. (B) Enrichment of dissimilar HRPs in human essential proteins. The grey histogram represents the random distribution and the pink arrow denotes the real observation. P-value and Z-scores were computed using permutation testing by performing 10,000 randomizations. (C) Bar plot showing the proportion of singleton HRPs and dissimilar HRPs that are essential in humans. n denotes the number of proteins in each class. P-value was estimated using Fisher’s exact test. Boxplot showing (D) the number of protein interactors and (E) the number of link communities that singleton and dissimilar HRPs participate in. n denotes the number of proteins in each class. P-value was computed using Wilcoxon rank sum test. (F) Schema depicting computation of change in interactability (Δinteractability) using divergent paralog pairs (top panel). Expected Δinteractability of multi-HR proteins was estimated as an exponentiation of the Δinteractability of singleton HRPs by the number of HRs in the multi-HR protein. Plot showing expected (line plot) and observed (bar plot) Δinteractability for proteins with dissimilar HR pairs. The numbers in the parenthesis denote the number of proteins harboring the corresponding number of HR(s). (G) Heatmap showing the fold change of observed probabilities of co-occurrence of dissimilar HR-pairs over expected probabilities (computed using singleton HRP estimates). P stands for probability.

### Dissimilar HRPs are physiologically important regulators of diverse spatio-temporal processes

Are dissimilar HRPs physiologically relevant? Our analysis shows that human essential repeat-containing proteins (E-HRPs) are significantly enriched in dissimilar HRPs [**Fig. 1B**], underscoring their importance for organismal survival. To further explore their functional roles, we compared dissimilar HRPs and singleton HRPs. Notably, dissimilar HRPs show a greater enrichment among human essential proteins than singleton HRPs [**Fig. 1C**]. Functional enrichment analyses revealed that dissimilar HRPs are predominantly associated with regulatory processes such as chromatin organization, transcription, as well as development and differentiation [**Supplementary Fig. 1D**]. Since both singleton and dissimilar HRPs participate in similar broad biological processes, we examined their specific molecular functions to identify distinctions. Interestingly, dissimilar HRPs are involved in different stages of histone-binding-related activities, suggesting specialized roles in epigenetic regulation, whereas singleton HRPs appear to function more generally in chromatin binding [**Supplementary Fig. 1E**]. Among development and differentiation related processes, dissimilar HRPs are associated with neuron fate commitment, neurogenesis and multi-organ development, while singleton HRPs are linked to vascular and urinary system development [**Supplementary Fig. 1F**]. Importantly, dissimilar HRPs exhibit greater developmental dynamicity, being dynamically expressed across a wider range of organs compared to singleton HRPs [**Supplementary Fig. 1G**]. This developmental dynamism reflects extensive variability in expression levels across different stages of development^22^. Such temporally dynamic expression patterns of dissimilar HRPs across multiple organs, could be a significant driver in shaping developmental outcomes. The ability of dissimilar HRPs to bring about global regulation in diverse biological contexts is best exemplified by (i) their participation in multiple regulatory phase-separated condensates involved primarily in transcriptional and post-transcriptional regulatory processes [**Supplementary Fig. 1H**] and (ii) broad tissue expression profiles compared to singleton HRPs [**Supplementary Fig. 1I**]. Overall, these findings suggest that proteins with dissimilar HRs could serve as key regulators of cellular functions across diverse spatial and temporal contexts.

### Protein interactability does not scale proportionally with the number of HRs

HRs are known to enhance the interactability of HRPs^15^, suggesting that an increase in the number of HRs within a protein may potentially lead to a proportional rise in its interaction partners. In line with this, we observed that dissimilar HRPs exhibit a higher number of protein interactors and protein-protein link communities compared to singleton HRPs [**Fig. 1D-1E**]. These findings suggest that an increase in the number and type of HRs within a protein enables (i) enhanced interactability and (ii) involvement in multiple protein complexes and/or pathways, as indicated by participation in more number of link communities. However, a closer examination of the distributions reveals only a modest increase in the number of interactions of dissimilar HRPs relative to singleton HRPs. In other words, the increase in the number of HRs does not result in a corresponding substantial shift in the distribution of the number of interactors. This suggests a possible interplay among different HRs within the same protein which may modulate the interaction potential. To investigate this, we analyzed human divergent paralog proteins, in which one paralog does not have HR (NonHRP), whereas the other protein is either a singleton HRP or a dissimilar HRP. We compared their interaction profiles while correcting for length-related biases. For this, we defined a parameter, Δinteractability, which quantifies the change in interaction profile associated with the acquisition of HR(s) [**Fig. 1F**; top schema]. Given that HRs are known to facilitate molecular interactions, one would expect that increase in the number of HRs would result in a proportional increase in Δinteractability [**Fig. 1F**; bottom schema]. To test this, we estimated the expected Δinteractability for dissimilar HRPs using the Δinteractability estimates of the singleton HRP divergent paralogs (null expectation) and compared this with the observed Δinteractability estimates for the dissimilar HRP divergent paralogs with different number of HRs. Surprisingly, we did not observe a proportional increase in the interactability of HRPs with acquisition of additional HRs [**Supplementary Fig. 2; Fig. 1F**]. This implies that multiple HRs within the same protein may not function independently. Instead, there appears to be an interplay between HRs that influences overall protein interactions. Collectively, our results indicate that different types of HRs coexisting within a protein may jointly modulate protein function.

### HR-mosaics are non-randomly distributed in the human proteome

If two distinct HR types within a protein (referred hereafter as HR-mosaics) function independently, the frequency of any given HR in a dissimilar HRP would be expected to mirror its frequency in singleton HRPs. To examine this, we first compared the distribution of HRs in dissimilar HRPs and singleton HRPs. We observed that certain amino acid HRs such as polyThr, polyGln and polyHis are more prevalent in dissimilar HRPs, whereas amino acids HRs such as polyVal, polyLeu and polyLys are underrepresented in dissimilar HRPs, relative to singleton HRPs [**Supplementary Fig. 3A**]. Does this distributional asymmetry reflect a tendency for specific HR-pair combinations to co-occur in dissimilar HRPs? To test this, we catalogued all distinct dissimilar HR-mosaics that co-occur within human dissimilar HRPs [**Supplementary Fig. 3B**], identifying each HR mosaic in the sequential order of their occurrence within the protein, while excluding similar HR-pairs.

The baseline probability of any given dissimilar HR mosaic co-occurring by chance in the human proteome (null expectation) was estimated using the singleton HR probabilities derived from the human proteins [**Supplementary Fig. 3C**]. Computation of fold change values for each dissimilar HR-mosaics as a ratio of the observed HR mosaic co-occurrence probabilities to the random expectation, enabled identification of the HR-mosaics that are enriched or depleted in the human proteome. Notably, HR-mosaics such as polyGly-polyHis, polyAla-polyHis, polyHis-polyGln show enrichment, whereas HR-mosaics such as polyGly-polyLeu, polyGly-polyLys, polyGlu-polyLeu show depletion [**Fig. 1G**]. These results are consistent with the previous findings that certain HR combinations occur at frequencies exceeding chance expectation^23^. Taken together these observations, point to (i) a preferential tendency for specific amino acid HRs to co-occur in dissimilar HRPs and (ii) non-random pattern of co-occurrence of dissimilar HR-mosaics in the human proteome.

### HR-mosaics can affect protein functionality through different modes of action

To quantify the extent of interplay between HRs in HR-mosaics, we developed a novel metric that estimates the magnitude of co-occurrence by using the instances of physical association of HRs within a mosaic in the same protein, across different evolutionary time-scales, ranging from days to millions of years. Physical association between any two HRs in the mosaics was estimated (i) within a cell represented by isoforms generated through alternative splicing, (ii) within a species, represented by gene duplicates or paralogs, and (iii) across species, represented by orthologs. We computed the HR physical association (HR-Pass) score for isoforms (HR-Pass_iso_), paralogs (HR-Pass_para_) and orthologs (HR-Pass_ortho_), and integrated these three estimates into an integrated Pass (HR-iPAss) score to distinguish mosaics with labile HRs (segregated mosaics) and those with HRs that tend to co-occur more often (concerted mosaics) [**Fig. 2A-2B**]. An HR-iPAss score closer to 0 (for instance, polyAsp-polyGlu and polyLys-polyGln mosaics) indicates that the HRs within a mosaic have minimal association across different time-scales. Conversely, a score near 1 (as seen for polyAsp-polyAla and polyLys-polyPro mosaics) signifies strong association of HRs within a mosaic. HR-mosaics with low HR-iPAss score indicate that each HR can act independently and contribute to distinct molecular outcome(s), which we define as segregated mode of action. In contrast, HR-mosaics with high HR-iPAss score indicate that both HRs influence the same molecular outcome, suggesting that both HRs are necessary for proper protein functionality, a pattern we define as concerted mode of action [**Fig. 2C**].

**Fig. 2.**
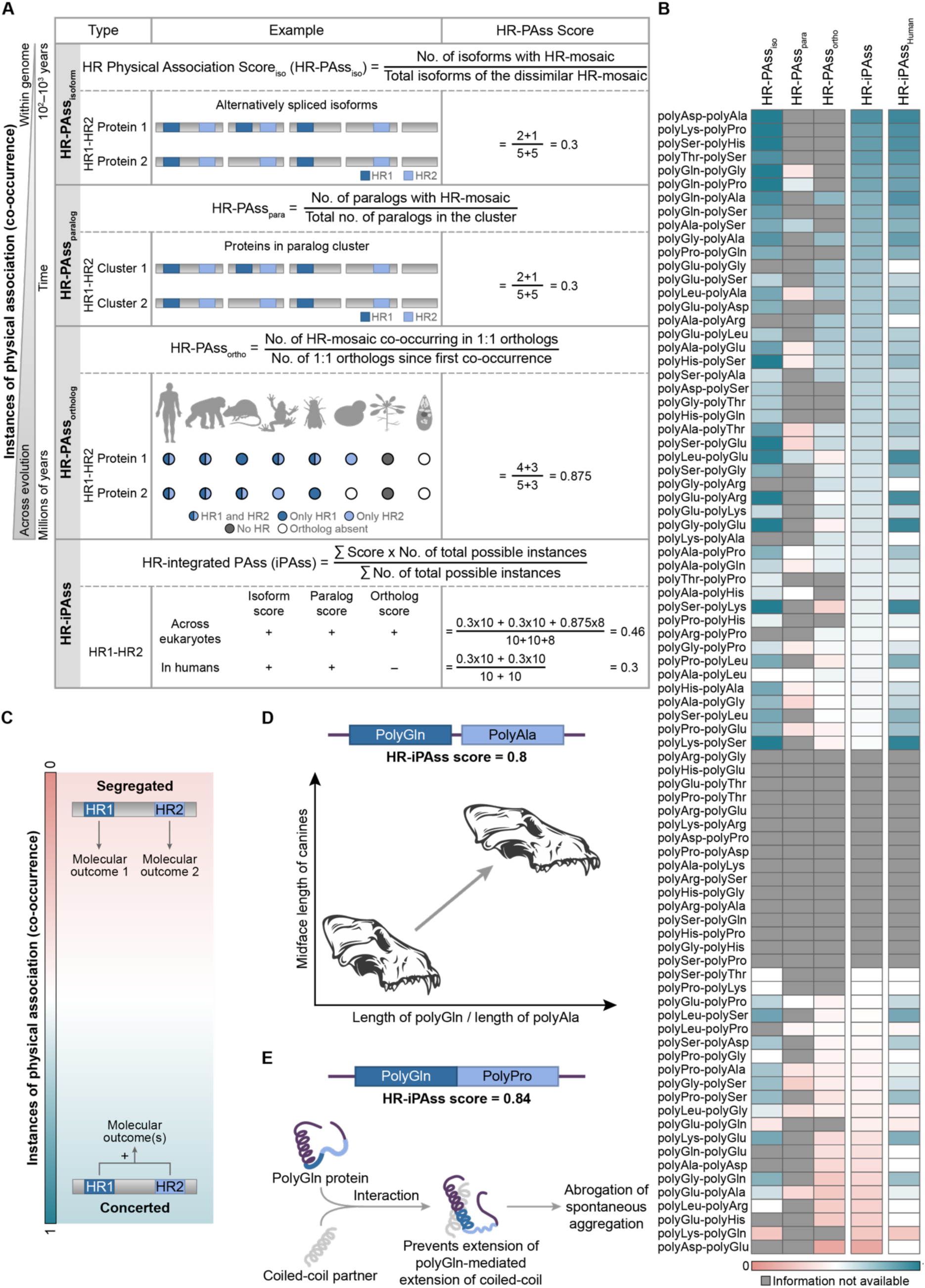
HR-iPAss scores delineate segregated and concerted HR-mosaics. **(A)** Computation of HR physical association (HR-PAss) score for isoforms, paralogs, orthologs and integrated HR-PAss (iPAss) score for dissimilar HRPs with two HRs. **(B)** Heatmap showing HR-Pass scores for different HR-mosaics for different physical association parameters. **(C)** Illustration showing segregated and concerted mode of actions of HR-mosaics. **(D)** Relative lengths of polyGln and polyAla HRs in RUNX2 show correlation with cranial skull morphology of canines^20,75^. **(E)** In chimeric peptides polyGln-polyPro HR-mosaic exhibits diminished spontaneous aggregation through coiled-coiled interactions upon encountering a coiled-coil containing binding partner^18,19^.

We hypothesized that concerted HR-mosaics may affect shared molecular outcome(s) through two distinct mechanisms: (i) a concerted disjunct mode, in which the HRs within a mosaic operate independent of each other, but collectively shape the molecular outcome(s), and (ii) a concerted conjunct mode, in which the HRs within a mosaic directly influence each other, and jointly contribute to the resulting molecular outcome(s). Crucially, existing literature lends support to our categorization of concerted-conjunct mosaics. For instance, polyGln-polyAla, with a high HR-iPAss score of 0.8, is classified as concerted conjunct mosaic [**Fig. 2D**]. In the Runt-related transcription factor 2 (RUNX2), which plays a key role in osteoblast differentiation, the relative lengths of polyGln and polyAla, has been associated with the midface length variation across dog breeds^20^. This aligns with the concerted-conjunct relationship between the two HRs identified here, implying that polyGln and polyAla may influence each other and jointly determine the phenotypic outcomes related to canine skull morphology. Similarly, in the polyGln-polyPro mosaic, polyPro on the C-terminus of polyGln can abrogate extended coiled-coil mediated spontaneous aggregation of polyGln when in contact with a coiled-coiled partner^18,19^, implying a concerted-conjunct relationship. Consistent with this, our analysis assigned the polyGln-polyPro mosaic an HR-iPAss score of 0.84 [**Fig. 2E**]. Based on these examples and given that HRs within concerted-conjunct mosaics can mutually influence one another, such mosaics would be expected to fall within the upper quarter of the HR-iPAss score distribution (approximately ≥0.75).

### PolyGly-polyPro mosaic in DDX17 shows concerted-disjunct mode of action

Does HR-iPAss score of around 0.5 reflect a concerted-disjunct relationship? To address this, we experimentally characterized the polyGly-polyPro mosaic of human DEAD box RNA helicase DDX17 [**Fig. 3A**] which was assigned an HR-iPAss score of 0.54. While the polyGly HR in DDX17 is conserved across eukaryotes, polyPro HR appears to have evolved in chordates [**Supplementary Fig. 4**]. To understand how the polyGly-polyPro mosaic affects DDX17 function, we generated DDX17WT, polyGly (DDX17ΔG), polyPro (DDX17ΔP) and the double (DDX17ΔGΔP) deletion mutants and transfected them into HEK293 cells [**Fig. 3A**]. We then assessed the impact of these HR deletions on protein abundance, subcellular localisation, protein mobility and protein-protein interactions. Regarding protein abundance, polyGly deletion resulted in a two-fold increase in steady-state levels of DDX17, whereas polyPro deletion led to a two-fold decrease [**Fig. 3B-3C**]. Notably, the double deletion (DDX17ΔGΔP) resulted in a modest change in the steady state levels relative to the DDX17WT, implying polyPro as the primary determinant of DDX17 abundance. In terms of subcellular localisation, DDX17WT was predominantly found in the nucleolus, as marked by fibrillarin [**Fig. 3D**]. While DDX17ΔG displayed a puncta-like distribution around the nucleolus with scattered nucleoplasmic puncta, DDX17ΔP displayed a nucleolar distribution albeit appearing diffuse. This could be partly attributed to the reduced abundance of the DDX17ΔP protein [**Fig. 3D**]. Notably, DDX17ΔGΔP double mutant exhibited localisation pattern comparable to DDX17WT suggesting a mutual buffering relationship between the HRs. Collectively, these observations suggest that polyGly is a key driver of nucleolar localisation of DDX17. We then assessed the influence of HRs on the mobility of DDX17 using fluorescence recovery after photobleaching (FRAP) experiments [**Fig. 3E; Supplementary movie 1-4**]. DDX17ΔG showed decreased mobility (referred as proteolethargy^24^), while DDX17ΔP displayed enhanced mobility (a phenomenon we term as proteolaxity) relative to DDX17WT. Remarkably, simultaneous deletions of both HRs restored the mobility to DDX17WT levels. Taken together, these results indicate that polyGly and polyPro tracts exert opposing and counterbalancing effects on DDX17 mobility.

**Fig. 3.**
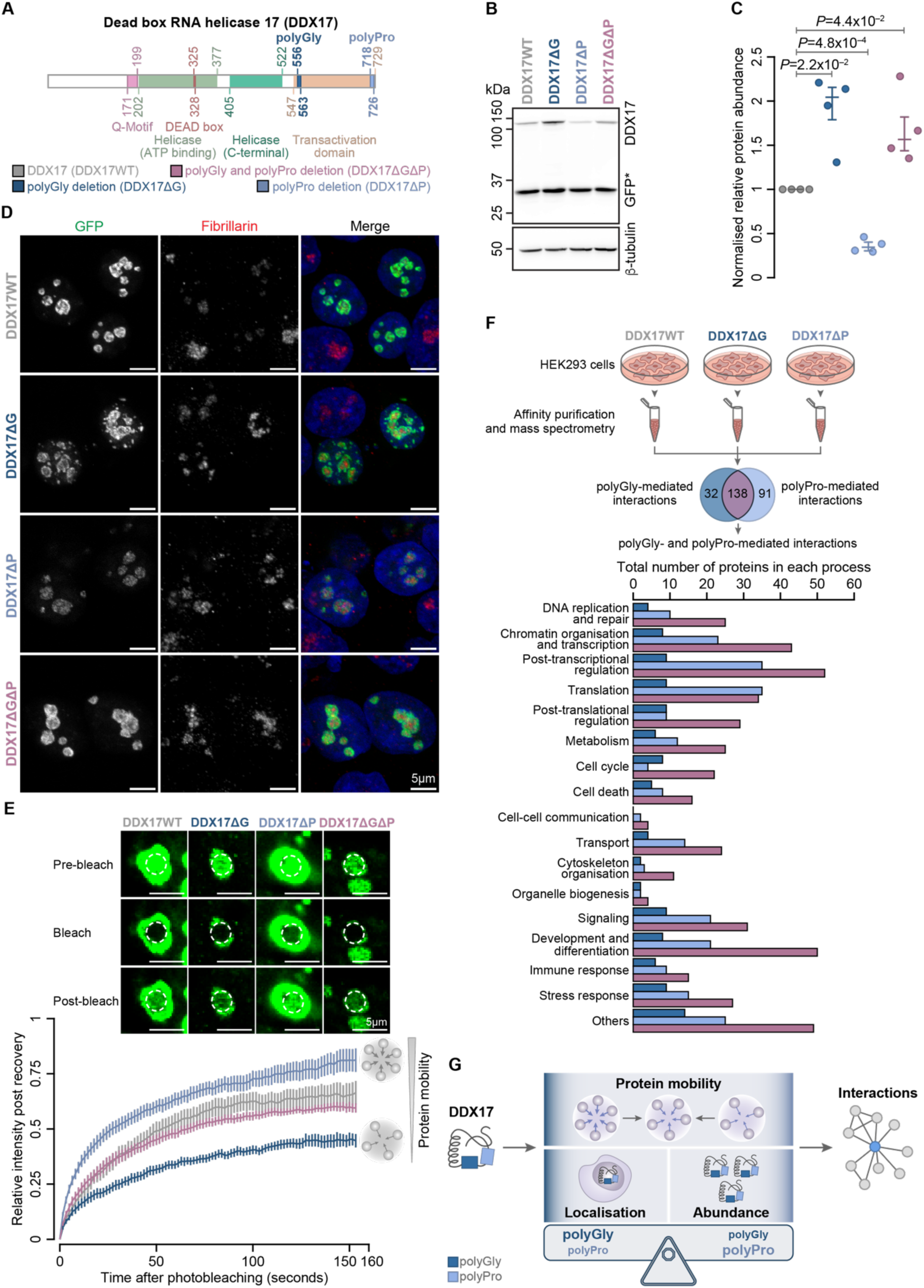
PolyGly-polyPro in human DDX17 exhibits concerted-disjunct mode of action. **(A)** Domain architecture of human DEAD-box RNA helicase 17 (DDX17), highlighting the locations of polyGly and polyPro HRs. **(B)** Western blot showing abundance levels of DDX17WT and DDX17ΔG, DDX17ΔP and DDX17ΔGΔP mutants (top panel). Asterix indicates empty GFP, which was used as a transfection control. Δ-tubulin is used as a loading control (bottom panel). **(C)** Quantification of the protein abundance of DDX17WT and DDX17ΔG, DDX17ΔP and DDX17ΔGΔP mutant proteins. Plot showing protein abundance relative to DDX17 WT as mean ± standard error of mean (SEM). P-value was estimated using Welch’s two-sample T-test. **(D)** Confocal images showing subcellular localisation of DDX17WT and DDX17ΔG, DDX17ΔP and DDX17ΔGΔP mutant proteins (GFP; green). Fibrillarin was used as a nucleolar marker (red) and nucleus was stained using DAPI (blue). **(E)** The top panel depicts the representative FRAP time course of GFP labelled DDX17 condensates of the indicated constructs. Dashed circles denote photobleached regions. Graph at the bottom shows FRAP recovery curves of DDX17 variants showing condensate mobility of different mutants in comparison to the DDX17WT. Data is presented as mean ± SEM. **(F)** Schema illustrating the workflow adopted to profile the protein interactors of DDX17WT and DDX17ΔG and DDX17ΔP mutant proteins using affinity-capture, followed by mass spectrometry (top panel). Bar plot showing the number of HR-mediated protein interactors of DDX17. Gene Ontology-Biological Processes (GO-BP) terms for each protein were obtained from QuickGO^38,39^ and manually classified into broad biological processes (bottom panel). **(G)** Illustration summarising the impact of polyGly and polyPro HRs on different molecular attributes of DDX17.

Considering the distinct molecular effects of the two HRs, especially on protein mobility, it is reasonable to expect that these repeats exert a substantial impact on the interaction landscape of DDX17. To explore this, we profiled the protein interactions of DDX17WT and the HR-deletion mutants using affinity capture mass spectrometry. We identified a total of 472 interactions of DDX17, of which 431 represent novel interactions, not previously reported [**Supplementary Fig. 5A**]. Strikingly, over 90% of these interactors were associated with distinct protein complexes, and more than 80% of them are constituents of phase-separated condensates, predominantly the nucleolus, stress granules and P-bodies [**Supplementary Fig. 5B-5C**]. These results are consistent with the established role of DDX17 as a multifunctional RNA helicase involved in the organization and regulation of ribonucleoprotein assemblies. We then categorized the interactions identified in DDX17WT but absent in different HR deletions as (i) polyG-mediated interactions, those that are exclusively lost in DDX17ΔG, (ii) polyp-mediated interactions, those lost exclusively in DDX17ΔP, and (iii) polyG-polyP-mediated interactions, those which are lost in both DDX17ΔG and DDX17ΔP [**Fig. 3F; Supplementary Fig. 5D**]. The largest subset of interactions required both polyGly and polyPro, followed by those that are selectively dependent on polyPro and then those that require polyGly. Gene Ontology Biological Process (GO-BP) annotations revealed that proteins requiring both polyPro and polyGly were enriched in all biological processes represented in the DDX17 interactome [**Fig. 3F**].

Collectively, these findings suggest that the two HRs in DDX17 serve distinct yet complementary roles wherein polyPro predominantly regulates protein abundance, polyGly principally drives localisation of DDX17 to the nucleolus and both HRs jointly determine the protein interaction landscape and thereby shape the overall functionality of DDX17 [**Fig. 3G**]. Thus, polyGly and polyPro in DDX17 influence different attributes of DDX17, they converge on the shared molecular outcome(s), namely determining the DDX17 interactome, without directly modulating each other. Thus, the polyGly-polyPro mosaic of human DDX17 exemplifies the concerted-disjunct mode of action.

### Relative positioning and length of HRs in a mosaic can shape the molecular outcomes

Does the position and length of the HRs within a mosaic impact protein function? To investigate this, HR-mosaics exhibiting significant correlations in the lengths of their constituent HRs across the human proteome were identified, yielding seven such HR-mosaics [**Fig. 4A**]. This length correlation revealed two surprising findings. First, polyGly-polyGln, which displayed length correlations in the human proteome, has been categorized as a segregated mosaic based on its HR-iPAss score (0.32). The length correlation between the two HRs was unexpected, since HRs within segregated mosaics are assumed to operate independently and influence distinct outcomes. This discrepancy prompted us to compute HR-iPAss_Human_ scores by integrating HR-PAss_iso_ and HR-PAss_para_ scores, excluding the macro-evolutionary relationships (HR-Pass_ortho_) [**Fig. 2B, 4A**]. This revised scoring reclassified polyGly-Gln as a concerted mosaic (HR-iPAss_Human_ score = 0.75), suggesting that the polyGly-polyGln pairing may have evolved to act synergistically in humans. Similarly, the HR-mosaics polyLys-polyGlu and polyPro-polySer, originally designated as segregated by the HR-iPAss score, were reassigned as concerted mosaics under the HR-iPAss_Human_ framework [**Fig. 2B**]. Collectively, these results indicate that the functional relationship between HRs within a mosaic may be species-specific, and that applying both integrated and species-specific scoring provides a more complete understanding of how HR-mosaics contribute to protein functionality.

**Figure 4.**
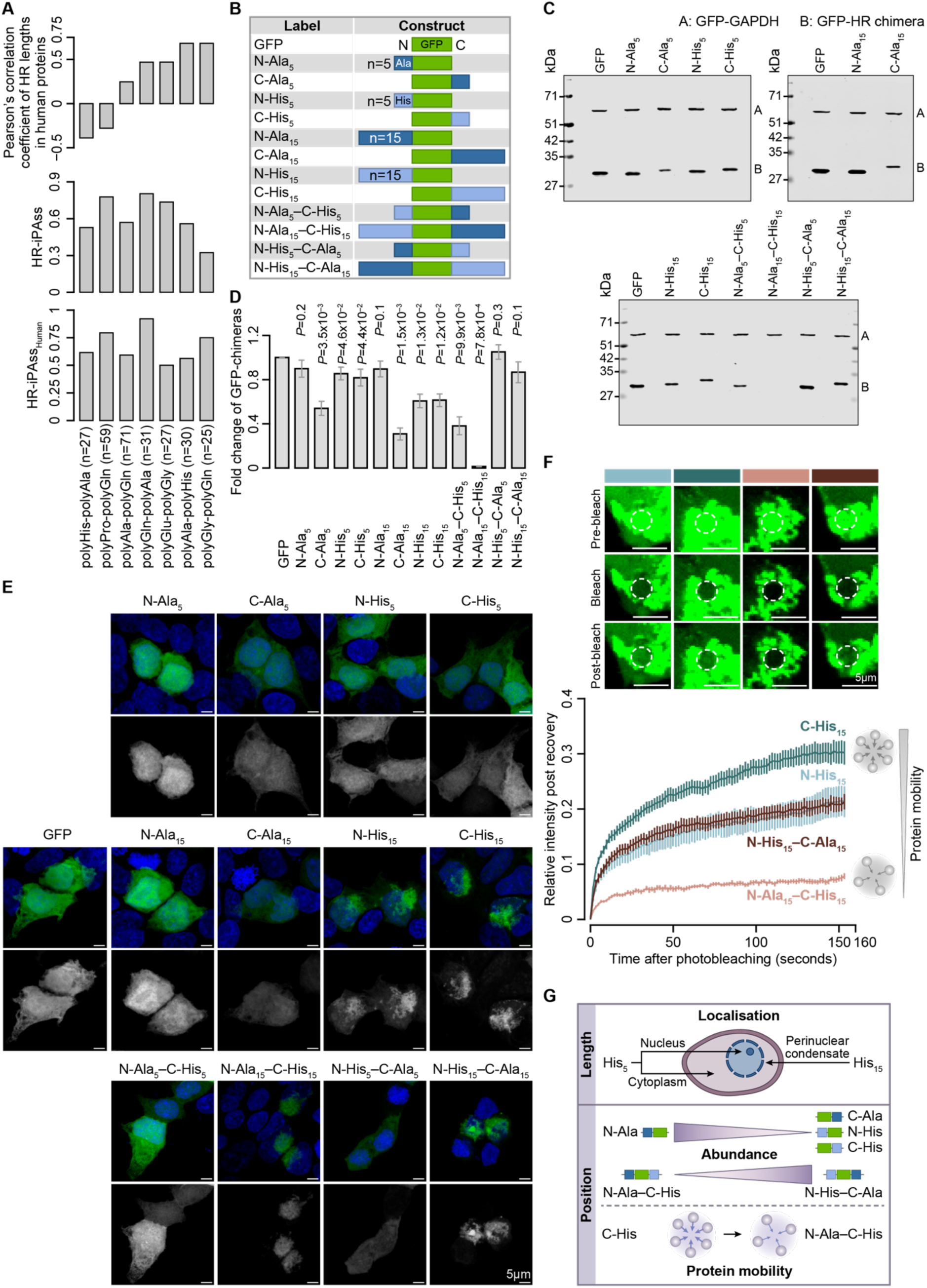
GFP chimeras with polyAla and polyHis show length- and/or position-dependent effects of HRs within HR-mosaics. **(A)** Bar plot showing HR pairs with a significant Pearson’s correlation of the length of the first HR with that of the second HR across human dissimilar (top), their HR-iPAss scores (middle) and HR-iPAss_Human_ scores (bottom). All dissimilar HRPs were used for computing correlation. HR-mosaics present in at least 5 dissimilar proteins were considered for the analysis. **(B)** A schematic representation of the various GFP chimeras generated in this study, encompassing individual or combined polyAla and polyHis repeats with differing lengths and positional arrangements. **(C)** Western blot showing abundance levels of the different GFP chimeras. **(D)** Plot showing protein abundance of GFP-chimeras as mean ± standard deviations. The levels of GFP or GFP-HR chimera proteins were normalized to the protein levels of GFP-GAPDH and presented as a relative value to the GFP protein. P-value was estimated using two-tailed T-test. **(E)** Confocal images showing the subcellular localisation of GFP chimeras (green). **(F)** Panel on top depicts representative FRAP time course of the indicated GFP chimeras. Dashed circles denote photobleached regions. Graph at the bottom shows FRAP recovery curves of the indicated GFP chimeras showing condensate mobility. Data is presented as mean ± SEM. **(G)** Illustration showing the influence of polyAla and polyHis HRs on different protein molecular attributes.

The second surprising finding was that two HR-mosaics sharing the same constituent HRs displayed opposing length correlation patterns depending on their relative positioning. Within the polyAla-polyHis mosaic, the lengths of polyAla and polyHis were positively correlated, whereas in the polyHis-polyAla mosaic, the same pair exhibited a negative correlation [**Fig. 4A**]. These contrasting trends imply that the functional contributions of HRs within a mosaic might be determined by their lengths and relative positioning. To investigate this, we constructed GFP chimeric proteins with varying lengths (5 or 15 amino acids), positions (N- or C-terminal relative to GFP) and combinations of polyAla and polyHis [**Fig. 4B**] and assessed their impact on protein abundance, localization and mobility. When HRs are individually present, (i) polyAla showed a marked position-dependent effect wherein the C-terminal HR dramatically reduced the steady-state protein levels compared to the effect at the N-terminus, while (ii) polyHis reduced protein abundance in a length-dependent manner, regardless of its position [**Fig. 4C-4D**]. When combined, C-terminal polyHis paired with N-terminal polyAla drove a pronounced reduction in abundance, indicating that C-terminal polyHis is sufficient to dictate steady-state levels. Conversely, while N-terminal polyHis and C-terminal polyAla each individually reduced protein levels, their combination surprisingly abolished this reduction. Furthermore, the longer polyHis (His_15_), promoted assembly or aggregate formation and altered intracellular localization irrespective of position, with these assemblies notably confined to the perinuclear region, unlike in the case of all other GFP chimeras, which distributed throughout the cell [**Fig. 4E**]. How does this affect the molecular outcomes when both HRs co-occur in mosaics with different relative positioning? FRAP analysis of constructs containing longer polyHis revealed that N-terminal His_15_ retained its mobility even when paired with C-terminal Ala_15_, whereas C-terminal His_15_ lost mobility upon pairing with N-terminal Ala_15_ [**Fig. 4F; Supplementary movie 5-8**]. Strikingly, constructs with identical repeat lengths but differing positional arrangements exhibited opposite recovery kinetics. As a relevant biological example, ZIC2 and ZIC3 proteins, members of the Zinc fingers in Cerebellum (ZIC) family of transcriptional regulators, that play essential roles in early embryogenesis and establishing body symmetry, harbor polyAla and polyHis stretches. This HR architecture might underpin precise regulation of their protein levels during early human development and its disruption could potentially compromise normal developmental trajectories or elevate the risk of cancer^25^. Collectively, these findings demonstrate that the biophysical properties of HR-containing assemblies are shaped by both the composition and positional arrangement of the constituent repeats [**Fig. 4G**].

### Functional code of HR-mosaics in humans

Is there a functional correspondence between HR-mosaics and distinct cellular processes? To examine this, Bayesian inference was employed by estimating conditional probabilities for proteins harboring HR-mosaics and the preferential association of dissimilar HR-mosaics with broad GO-biological processes was examined [**Fig. 5A**]. This analysis identified particular HR-mosaics that are predominantly associated with specific biological processes [**Fig. 5B**]. Notably, HR-mosaics in proteins associated with nucleic acid binding processes such as DNA replication and repair, chromatin organisation, transcription and translation show a tendency for concerted mode of action. Conversely, processes involved in cell-cell communication are predominantly associated with HR-mosaics that exhibit a segregated mode of action. Jaccard Similarity Index (JSI) estimations between different biological processes revealed considerable overlap of HR-mosaics which predominantly operate in a concerted manner across processes like chromatin organisation and transcription, post-translational regulation, signaling as well as development and differentiation [**Supplementary Fig. 6A**]. On the other hand, processes such as translation and organelle biogenesis share limited HR mosaics with other biological processes, suggesting that specific HR mosaics may be engaged in these processes.

**Fig. 5.**
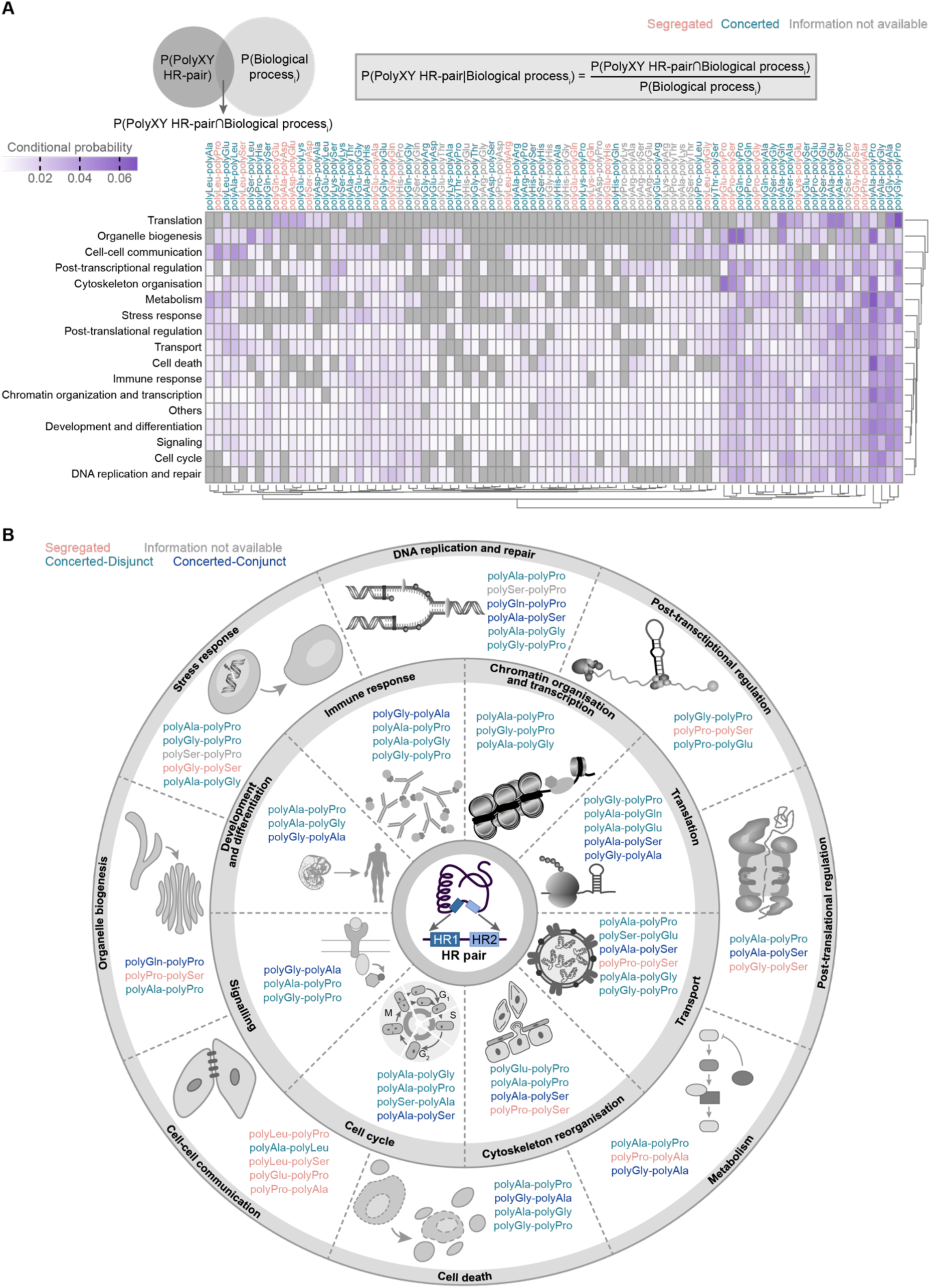
HR-mosaics are preferentially associated with distinct biological processes. **(A)** Heatmap depicting the conditional probability estimates of dissimilar HR pairs involved in different biological processes. Gene ontology biological processes (GO-BP) annotations were mapped for each dissimilar HRP from QuickGO^38,39^ and manually classified into broad biological processes. **(B)** Illustration depicting the top HR pairs with highest conditional probabilities that are preferentially associated with different biological processes. The top three HR-mosaics associated with each biological process, based on conditional probabilities, are represented in the figure.

Analysis of proteomes from various phase-separated condensates revealed that HR-mosaics operating in segregated mode such as polyGly-polySer is present in as many as 11 distinct condensates [**Supplementary Fig. 6B**]. This broad participation may reflect an expansion of functional space, whereby each HR acts independently to drive multiple distinct molecular outcomes. In contrast, concerted-conjunct HR-mosaics such as polySer-polyHis, polyGln-polyAla and polyGln-polySer, are found in a comparatively limited number of phase-separated condensates. This restricted distribution may suggest that the mutual interdependence between HRs in these mosaics has driven their evolutionary specialization toward participation in specific condensates, enabling the execution of defined functional outcomes. These results highlight the presence of pleiotropic and process-specific HR-mosaics. Collectively, these findings establish the existence of distinct functional HR-mosaic codes, characterized by distinct modes of action and preferential association with distinct cellular processes in humans.

## Discussion

Prior studies investigating the molecular roles of individual HRs have established them as important functional units within disordered protein regions^13–17,25–34^, yet how combinations of HRs, which we term as HR-mosaics, collectively influence protein functionality remains largely unexplored. We present a comprehensive systems-level analysis of HR-mosaics comprising of different HR types within the same protein and their collective influence on protein function(s). To achieve this, we developed a novel evolutionary co-occurrence metric, which enabled the categorization of HR-mosaics by their modes of action in both evolutionarily conserved and species-specific contexts. The molecular case studies presented here, alongside the literature-supported examples affirm the utility of this theoretical framework to characterize the functional interplay between co-occurring HRs and the extendibility of this framework across species. These findings suggest that evolutionary pressures may have shaped the specific combinations of amino acid HR types, their lengths and their relative positioning in accordance with the biological processes and compartments (both membrane-bound and membrane-less) to achieve specific biochemical outputs [**Fig. 6A**]. This is exemplified by the non-random organizational patterns of HRs within mosaics and the existence of distinct functional HR-mosaic codes associated with specific biological processes.

**Fig. 6.**
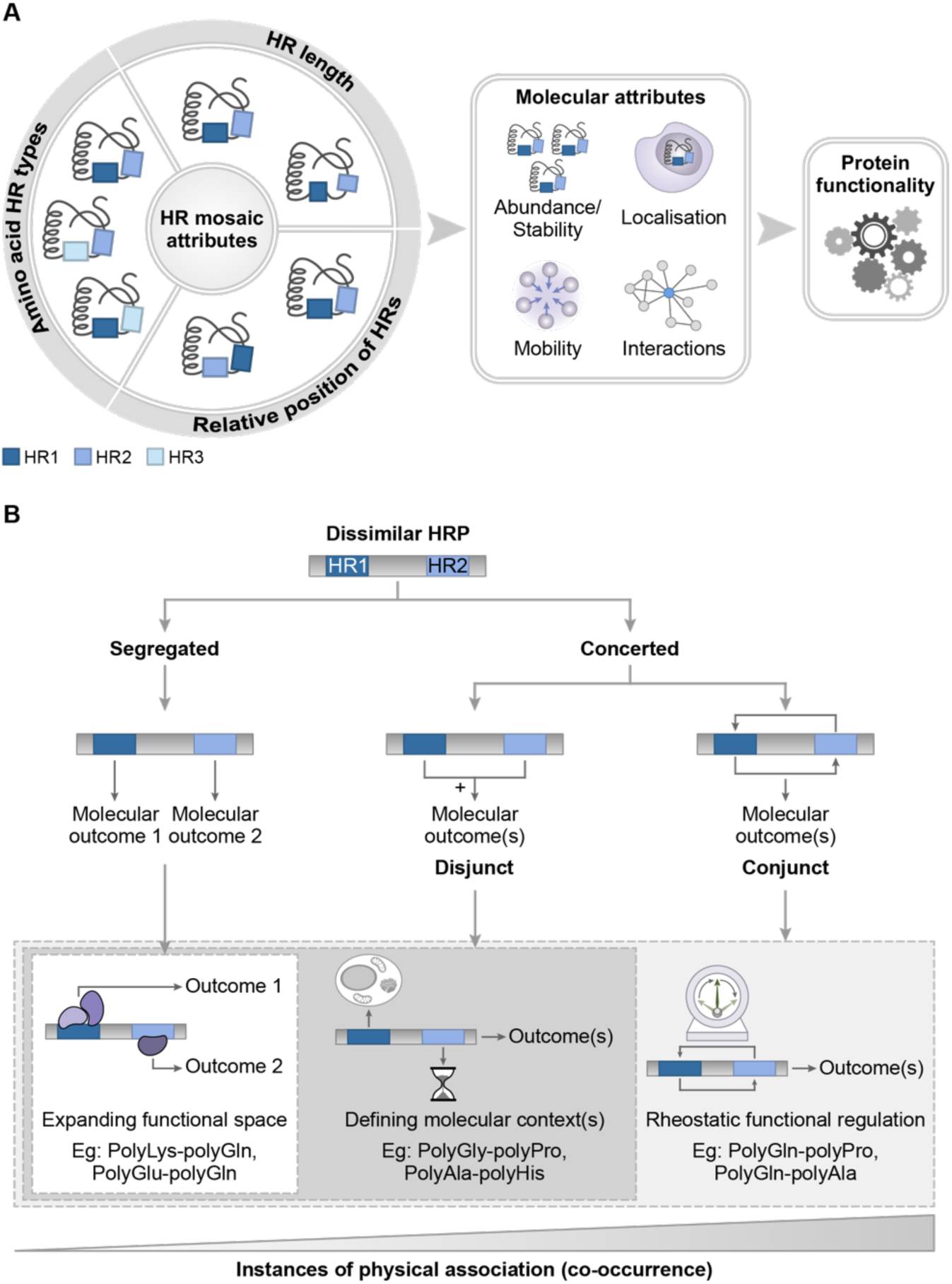
The influence of HR-mosaics on the protein functional landscape. **(A)** Key attributes of HR-mosaics, including the identity of the constituent amino acid HRs, their lengths and their relative positioning, collectively orchestrate protein functionality by modulating protein abundance, subcellular localization, molecular mobility and interaction networks. **(B)** A schematic representation of the distinct modes of action adopted by HR-mosaics and their corresponding functional manifestations.

The various modes of action through which HR-mosaics operate, contribute to distinct functional facets of protein behaviour. For instance, when two HRs within a protein operate in a segregated mode, each independently governs distinct molecular outcomes, collectively broadening the functional repertoire of the protein. This expansion of functional space may underlie the enhanced interactability and greater participation in multiple phase-separated condensates seen in proteins harboring dissimilar HR mosaics. In the concerted-disjunct mode, the HR-mosaics not only broaden the functional space but also confer molecular context to the protein, such as defining its spatiotemporal localization or modulating its abundance. In the concerted-conjunct mode, through direct mutual influence, the two HRs additionally enable rheostatic regulation of protein functionality, generating a continuum of functional outputs [**Fig. 6B**]. The interdependence inherent to co-occurring HRs, thus presents opportunities not only for accessing wider functional landscape but also for precise fine-tuning of protein functionality, achievable by modulating HR lengths or altering the spatial distance between them. In fact, some of the amino acid HRs are longer in the mosaic configuration [**Supplementary Fig. 7A**], than when exist alone, which we term as SupraHR in line with the Supradomains found in the structural regions of proteins^6^. Although this study primarily examined proteins harboring two dissimilar HRs, many proteins contain multiple distinct HR types, presenting an enormous combinatorial potential. This implies that within a single protein, several HR-mosaics with different modes of action may operate simultaneously, each contributing to distinct functional dimensions. The functional logic governing individual HRs and their combinations is non-linear, wherein the outputs can vary considerably in magnitude and, in certain instances, shift to completely opposing outcomes [**Fig. 4G**] highlighting the complexity and richness of HR-mosaic encoded functionality.

Although human dissimilar HRPs are predominantly conserved across eukaryotes [**Supplementary Fig. 7B**], they display a greater degree of repeat-associated variation compared to singleton HRPs [**Supplementary Fig. 7C**]. This suggests that at the population level, HR-mosaics harboring variable HRs may drive both discrete and continuous functional outputs. The standing genetic variation arising from such repeat-associated variability could provide populations with ready access to a broad adaptive potential, enabling rapid and robust responses to sudden environmental challenges. More importantly, in the context of HR-mosaics, minor changes in HR length or relative positioning can function as molecular leverage points that precipitate into huge shifts in biochemical outputs, particularly given that HR-mosaics are enriched in global regulators endowed with expansive regulatory potential.

Once dismissed as functionally inert, the findings presented here demonstrate that HRs individually or in combination can encode multiple functions, influencing several determinants of protein function. We thus establish these repeats as functionally versatile units capable of acting as stability codes affecting protein turn-over, localization signals for different sub-cellular compartments^30^, distribution codes for different chemical ^35^, mobility codes that determine protein dynamics^24^, and interaction meidators^15,25–27^, and propose the term “Versons” to describe them.

This study further establishes that appropriate protein mobility can be maintained through the counterbalancing effects of two distinct HRs. In the context of repeat-expansion diseases, this implies that the molecular etiology should be examined through the lens of HR-mosaics as a whole, rather than focusing solely on the disease-associated repeat in isolation. Notably, a substantial proportion of HR-expansion diseases are linked to dissimilar HRPs [**Supplementary Datasheet 1**], and that proteolaxity, analogous to proteolethargy, may itself contribute to molecular pathogenesis. Even in the context of infectious disease, dissimilar HRPs appear to be preferentially engaged in human-pathogen protein-protein interactions [**Supplementary Fig. 7D**], further underscoring their biological significance. Looking ahead, large-scale investigations encompassing diverse HR-mosaic combinations, probing multiple functional outputs across varied cellular environments, and leveraging machine and deep learning approaches are anticipated to yield a comprehensive understanding of HR-mosaic logic and how these principles can be harnessed for rational protein design and applied toward synthetic and translational objectives.

## Methods

### Identification of amino acid homorepeats in the human proteome

We obtained the sequences of all human proteins from the UniProt SwissProt database. We used an in-house python script to identify homorepeats (HR) in the human proteome. A protein with at least one HR stretch reaching a minimum of 5 residues was classified as an HRP, as previously defined^15^. Proteins with only one HR were classified as singleton HRPs, proteins with more than one HR of the same amino acid type were classified as similar HRPs and those with more than one HR of which at least one was of a different amino acid type were classified as dissimilar HRPs [**Supplementary datasheet 1**].

### Gene Ontology related analysis

We estimated the enrichment of Gene Ontology-Biological Processes (GO-BP) DAVID server^36,37^. An FDR <0.05 was used to estimate significance. The enriched GO-BP terms were then manually categorised into major biological processes and functions. For each dissimilar HRP, GO-BP was obtained from QuickGO^38,39^ and each term was manually categorised into major biological processes [**Supplementary datasheet 1**].

### Assembling human proteome-level datasets

Human essential genes/proteins were collated from different mid- to high-throughput studies^40–50^. Experimentally annotated human physical protein-protein interaction network was assembled from BioPlex^51^, CORUM^52^, DIP^53^, HINT^54^, PathwayCommons^55^, HuRI^56–61^ and BIOGRID (physical interactions not identified using Yeast2Hybrid system)^62^ as well as high-throughput studies^60^. The dataset for human proteins that undergo phase separation was assembled from different databases such as DrLLPS^63^, LLPSDB^64^, PhaSepDB^65^ and PhasePro^66^. We further assigned each condensate to different biological functions using extensive literature search. Tissue-specific expression was obtained from Human Protein Atlas^67^. Human developmentally dynamic genes/proteins were assembled for seven different organs^68^. Human paralog pairs were obtained and curated from Dandage et al^69^. Gene ages of human proteins were obtained from Singh AK et al^26^. Dataset for HR variability was obtained from Mier et al^70^. Human proteins involved in human-pathogen protein interactions were obtained from Singh et al^27^.

### Network Analysis

We estimated the topological property of connectedness between proteins (degree) in protein-protein interaction network using Cytoscape 3.0 and/or R. We identified the link communities of each protein in the human protein-protein interaction network using a previously described algorithm^71^.

### Estimation of HR-physical association scores

We assembled protein isoforms of human dissimilar HRPs and their sequences from UniProt SwissProt database, and identified the HRs using an in-house Python script, and subsequently classified them into singleton, similar and dissimilar HRPs. To compute the HR physical association score for protein isoforms (HR-PAss_iso_) with a HR pair (designated as HR1-HR2), we assembled isoforms of all dissimilar proteins with HR1-HR2 pair and computed the ratio of the total number of isoforms in the human proteome that contain the HR1-HR2 pair over the total number of all isoforms of all dissimilar proteins with that particular pair. The score was computed for HR pairs which have at least 5 isoforms for all dissimilar HRPs in the human proteome with the HR pair.

We collated paralogs of dissimilar HRPs^69^ and computed the HR-PAss score for paralogs (HR-PAss_para_) with any HR pair by computing the ratio of the total number of paralogs that have the HR1-HR2 pair over the total number of paralogs of all dissimilar proteins with HR1-HR2 pair. The score was computed for HR pairs which have at least 5 proteins in paralog clusters for all dissimilar HRPs with HR1-HR2 pair. We acquired the orthologs of human dissimilar HRPs across 460 different eukaryotic species from OMA browser^72^ and curated it to preserve only 1:1 orthologs across other species. Only one strain per species (strain with highest number of proteins or widely studied model strain) was selected as the representative. The species were then classified into their taxonomic lineages based on classification obtained from NCBI taxonomy browser^73^ and manual curation. To compute the HR-PAss score for protein orthologs (HR-PAss_ortho_) with an HR pair, we assembled orthologs of all dissimilar HRPs with the HR pair in the same order and nature of HRs as in humans. We then computed the ratio of the total number of protein orthologs that have HR1-HR2 pair over the total number of protein orthologs of all dissimilar proteins with HR1-HR2 pair since the first co-occurrence of HR1-HR2 ortholog was observed. We considered HR pairs which have at least a total of 5 orthologs identified for all dissimilar HRPs with HR1-HR2 pair. We then computed an integrated HR-PAss (HR-iPAss) score using the weighted average of HR-PAss scores across isoforms, paralogs and orthologs. For an HR pair with any missing value for any HR-PAss parameter, the specific HR-PAss score was replaced with 0.5 (keeping the n value 5) for computing the HR-iPAss score. All the scores are provided for each dissimilar HR mosaic in **Supplementary datasheet 1.**

### Cloning and generation of DDX17 HR-deletions

Wild-type DDX17 (DDX17WT) full-length corresponding to p82 isoform (NM_006386.5) was cloned from HeLa cDNA into pEGFPC1 vector backbone (Clontech) using EcoR1 and BamH1 sites. The constructs containing polyGly (DDX17ΔG) and polyPro (DDX17ΔP) deletions were synthesized in the pUC57 vector with BamHI and KpnI overhangs. Since the polyPro tract was at the end of the DDX17 coding region, the subsequent 3 amino acids after polyPro were also deleted. For the double deletion of the polyGly and polyPro tracts (DDX17ΔGΔP), GFP-DDX17ΔP was used as the template and deletion PCR was performed using the Q5 Site-Directed Mutagenesis Kit (E0554S, NEB), using primer mentioned in **Supplementary Table 1**. All constructs were verified by DNA (Sanger) sequencing.

### Cell culture and transfection for DDX17 HR-deletions and GFP-chimeras

HEK293 cells (ATCC) used for all the experiments were grown in DMEM GlutaMAX (10569044, Gibco) supplemented with 10% FBS (A55256701, Gibco) under 37°C and 5% CO2 incubation in a humidified chamber. Cultures were frequently tested for mycoplasma using PCR-based methods. The overexpression constructs were transfected using Lipofectamine 2000 transfection reagent (11668027, Invitrogen) according to the manufacturer’s protocol.

### Immunostaining of DDX17 HR-deletions and GFP-chimeras

Cells grown on coverslip/ chamber slide (177402, Thermo) were transfected and fixed after 36 hours with Image-iT Fixative 4% Paraformaldehyde (I28800, Thermo) for 20 minutes. Cells were permeabilised with 0.5% Triton X-100 in PBS for 20 minutes and blocked with 5% BSA in PBST containing 0.5% Tween 20 for 1 hour. Subsequently, cells were incubated with a primary antibody in blocking buffer at 4°C overnight, and a corresponding fluorescent secondary antibody for 1 hour at room temperature. The cells were washed 3 times after primary and secondary antibody incubation with PBST, then incubated with DAPI for 10 minutes before mounting with Prolong Gold antifade mountant (P36930, Invitrogen). Antibodies used are Fibrillarin Monoclonal Antibody (MA3-16771, Invitrogen) 1:300 dilution and Anti-mouse IgG Alexa Fluor 555 (A-21422, Invitrogen) 1:500 dilution. Images were acquired in FV3000 laser scanning confocal microscope (Olympus) under 60X objective and analysed in ImageJ^74^.

### Immunoblot analysis of DDX17 HR-deletions

HEK293 cells were co-transfected with DDX17 constructs along with empty pEGFPC1 vector and lysed after 36 hours in RIPA buffer containing protease inhibitor cocktail (11836170001, Roche). The soluble protein was quantified using BCA assay (T9300A, Takara) and equal protein were resolved on a 10% SDS PAGE gel under denaturing conditions. The resolved proteins were transferred onto a nitrocellulose membrane and were checked using Ponceau-S staining. The membrane was blocked in 5% non-fat dry milk (SC-2324, Santa-cruz) in TBST with 0.5% Tween20. The primary antibody was probed overnight at 4°C under constant rocking and the secondary antibody was probed for 1 hour at room temperature. The membrane was washed 3 times with TBST containing 0.5% Tween 20 after both primary and secondary antibody incubations. Antibodies used are: GFP (D5.1) antibody (2956, CST) 1:2000 dilution; Beta-tubulin antibody (2146, CST) 1:2000 dilution; HRP linked anti-rabbit IgG (7074, CST) 1:3000 dilution. The blots were imaged using chemiluminescence (1705062, Biorad) in Amersham AI800 imager (Cytiva) and quantified through densitometry in ImageJ^74^. The relative protein levels of DDX17 constructs were normalised to empty GFP co-expressed.

### Immunoprecipitation and mass spectrometry

Immunoprecipitation of DDX17WT, DDX17ΔG and DDX17ΔP was carried out using GFP trap beads (gta, Chromotek) as per the manufacturer’s instructions. Briefly transfected HEK293T cells were lysed using RIPA buffer, and GFP-tagged constructs of interest were immunoprecipitated using GFP-trap nanobeads as per the manufacturer’s protocol and precleared with agarose beads for 30 minutes. Equal amount of total protein (∼700 mg) was diluted five-fold in lysis buffer minus triton-X-100 and 20 ml of equilibrated GFP trap beads were added. The incubation was carried out at 4°C with rotation for 4 hours and then washed 3X with lysis buffer and twice with dilution buffer for 2 minutes each. Subsequently, samples were lysed in 2X SDS sample buffer at 80°C for 10 minutes. Quality of IP was assessed using immunoblot analysis. Subsequently, this immunoprecipitate was run on a precast SDS-PAGE gel and the gels bands were excised. The gel pieces were then washed, reduced, alkylated and digested using trypsin to generate peptides. The peptides were extracted from the gel by shaking in formic acid and acetonitrile followed by sonication. The samples were desalted using C10 Zip-Tip and processed for mass spectrometry. The raw mass spectrometric data was processed and analysed using Proteome Discoverer 1.4 with full human proteome (Jan 2020; UniProt) as the target database. We filtered the search results at 1% FDR at the peptide confidence level and at least two peptides per protein at the protein identification level. Proteins identified as interactors of DDX17WT were identified by removing protein interactors identified for GFP only. To identify interactors dependent on polyGly and polyPro HRs, we identified the protein interactors of DDX17WT (pEGFP-DDX17WT) that were not identified as interactors of DDX17ΔG (pEGFP-DDX17ΔG) and DDX17ΔP (pEGFP-DDX17ΔP), respectively.

### Fluorescence recovery after photobleaching (FRAP) of DDX17 HR-deletions and GFP-chimeras

FRAP was performed in live cells grown on chambered cover glass (155379, Thermo) after 36 hours of transfection. The cells were maintained in phenol-red-free media during the imaging, and FRAP acquisition was performed in FV3000 laser-scanning confocal microscope (Olympus). Following a few pre-bleach scans, a specific circular zone was marked for GFP bleaching with 100% laser. Fluorescent recovery was then continuously captured for about 2 minutes. The FRAP data was analysed in Olympus CellSens software to construct recovery curves in a single-exponential fit considering for the background and bleach corrections. FRAP time course of each protein is provided as **Supplementary movie 1-8**.

### Generation of GFP-chimeric proteins

To investigate the effect of Ala and His repeats on the properties of protein, GFP expression constructs were generated in which the GFP is tagged with Ala and/or His repeats of variable lengths at the N- or C-terminal ends of GFP. For this, the sequence encoding EGFP were PCR amplified using the pEGFPC1 vector (Clontech) with the primers listed in **Supplementary Table 2** and was cloned into the pCDNA4-myc/his vector (Invitrogen) using BamHI and XhoI restriction sites to generate pCDNA4-GFP and pCDNA4-GFP-repeat(s) constructs. The desired Ala or His repeats were incorporated in the PCR primers. The sequence encoding the human GAPDH was PCR amplified using the cDNA prepared from HEK293 cells and cloned into the pEGFPC1 vector to generate the pEGFP-GAPDH construct.

### Protein stability investigations of GFP-chimeras

To study the effect of His or Ala repeats on protein stability, 293T cells were co-transfected with pEGFP-GAPDH (transfection control) and pCDNA4-GFP or pCDNA4-GFP-repeat(s) construct at a ratio of 2:1. After 48h of transfection, the transfected cells were harvested and lysed and the lysates were subjected to immunoblotting with anti-GFP antibody (sc-9996, Clontech; 1:2000 dilution) and IRDye® 800CW secondary anti-mouse antibody (926-32210, LICORbio; 1:20000 dilution). All the western blot images were acquired using Odyssey DLx NIR fluorescence imager (LICORbio) and band intensities were quantified using ImageJ software across the samples. The levels of GFP or GFP-HR chimeric protein signals were normalized to the protein levels of GFP-GAPDH and presented as a relative value to the GFP protein.

## Quantification and statistical analysis

Statistical significance for differences of the distribution of discrete variables and continuous variables was estimated using Chi-squared or Fisher’s exact test, and the non-parametric Wilcoxon rank sum test, respectively. We corrected for multiple hypothesis testing using the Benjamini–Hochberg (FDR) method. Enrichment of proteins in different classes (for instance, dissimilar HRPs in proteins participating in different phase-separated condensates) was estimated by permutation testing performed through 10,000 randomizations. Every permutation involved substituting a random protein for each protein of interest (such as proteins involved in a particular phase-separated condensate). The number of random proteins that overlapped with a particular class of proteins (such as dissimilar HRPs) was recorded for every randomisation. We computed Z-score and P-value by comparing the random expectation and the actual observation. Z-score reflects the level of departure of the actual observation compared to the mean of random distribution in terms of the number of standard deviations. We evaluated P-values for enrichment as the ratios of the number of the randomly observed proteins ≥ the number of actually observed proteins to the total number of randomized samples (10,000). We computed conditional probabilities for finding a dissimilar HRP with polyHR1-polyHR2 mosaic in a specific biological process to obtain Bayesian inferences. All statistical analyses were performed in R.

## Supporting information

Supplementary Datasheet 1

## Acknowledgements

This study was supported by IISER Tirupati core funding (S.G., P.L.C., and S.C.); Prime Minister’s Research Fellowship, Government of India (A.K.S., N.R.); Ramalingaswami Re-entry Fellowship BT/RLF/Re-entry/05/2018 from Department of Biotechnology, Government of India (S.C.); Junior/Senior Research Fellowships from Department of Biotechnology, Government of India (K.S.K.); Department of Science and Technology INSPIRE fellowship (V.S.); Junior/Senior Research Fellowships from University Grants Commission, Government of India (A.S.G., A.B); Core Research Grant CRG/2023/004691 from Anusandhan National Research Foundation, Government of India (A.K.S., and S.C.); SERB Power grant SPG/2022/001987 (P.L.C.) and Wellcome trust India Allliance Intermediate grant IA/2019/I/1/504280 (P.L.C.).

## Author contributions

Conceptualization: S.C., P.L.C., A.D., A.K.S.; Methodology: A.K.S, A.S.G., P.L.C., A.D., S.C.; Investigation: A.K.S., A.B., A.S.G., V.S., S.G., K.S.K., H.R., N.R., P.S.; Visualization: A.K.S., S.C., P.L.C., A.D., A.B., A.S.G., V.S., K.S.K., N.R.; Funding acquisition: S.C., P.L.C., A.D.; Project administration: S.C.; Writing– original draft: S.C., A.K.S., P.L.C., A.D.; Writing–review & editing: A.K.S., A.B., A.S.G., V.S., K.S.K., N.R., A.D., P.L.C., S.C. All authors have read and approved the manuscript.

