## Supplementary Information for "Amino acid repeat mosaics shape protein functional landscape"

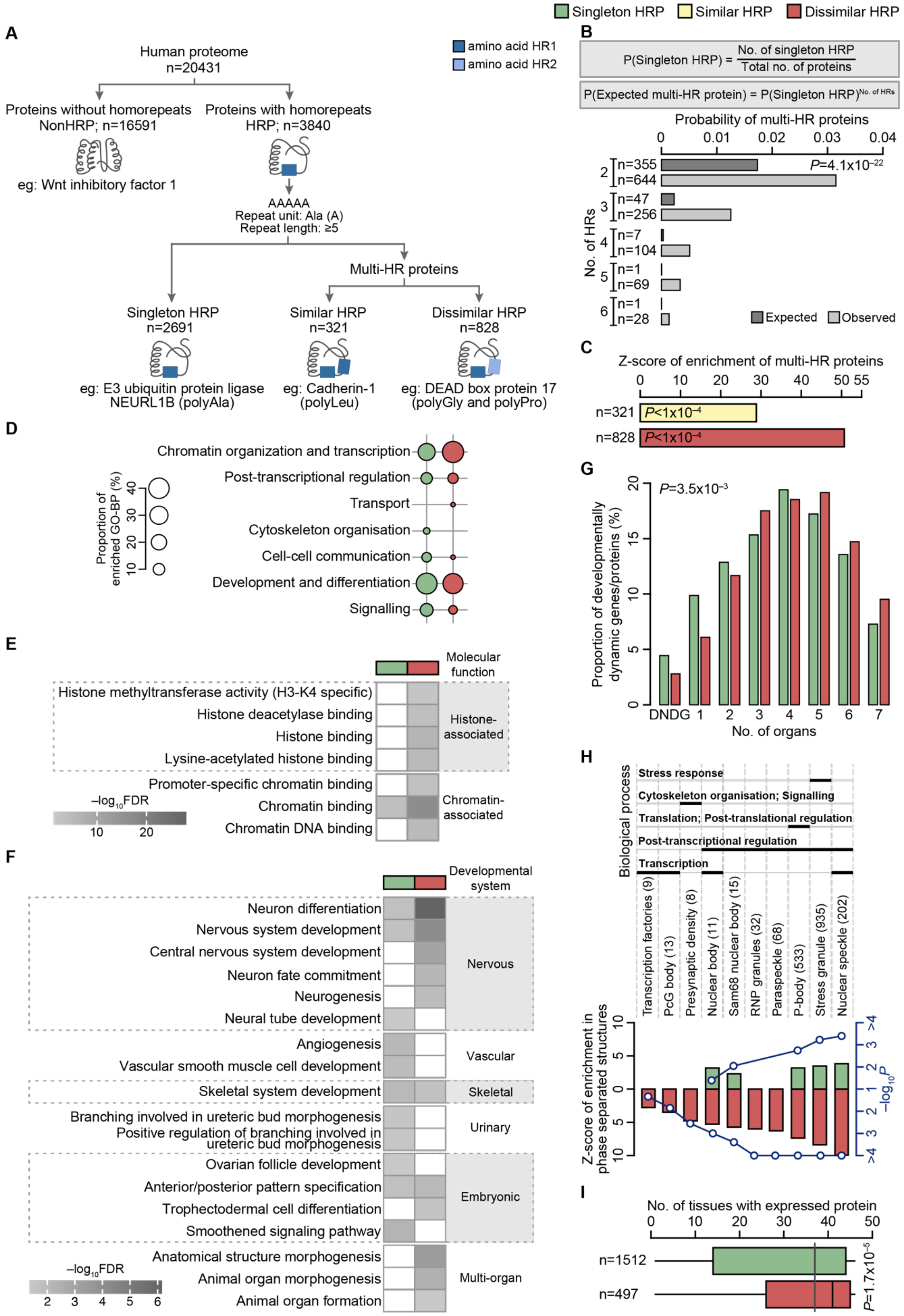


**Supplementary Fig. 1. Dissimilar HRPs are physiologically important. (A)** Classification of the human proteome based on the presence, number and type of HRs in each protein. **(B)**

Bar plot depicting the probability of multi-HR proteins in the human proteome. P-value computed using Chi-squared test. The expected probability for a multi-HR protein was estimated by exponentiation of the probability of observing singleton HRP in the human proteome by the number of HRs found in that particular multi-HR protein. **(C)** Bar plot showing the enrichment of similar and dissimilar HRPs in multi-HR proteins assessed using permutation testing by performing 10,000 randomizations. n denotes the number of proteins in each category. **(D)** Bubble plot showing significantly enriched (FDR<0.05) Gene ontology biological process (GO-BP) terms categorized as broad biological processes for singleton and dissimilar HRPs. **(E)** Heatmap showing significantly enriched (FDR<0.05) Gene ontology molecular function (GO-MF) terms categorized under histone- and chromatin-associated functions for singleton and dissimilar HRPs. **(F)** Heatmap showing significantly enriched (FDR<0.05) GO-BP terms categorized under development and differentiation for singleton and dissimilar HRPs. **(G)** Proportion of developmentally dynamic genes (DDGs) among singleton and dissimilar HRPs in seven different organs. P-value was computed using Fisher's exact test. DNDG represents developmentally non-dynamic genes. **(H)** Enrichment of singleton and dissimilar HRPs in different phase-separated condensates. The number in parenthesis indicates the total number of proteins in the condensate. Condensates with at least 10 proteins were selected for the analysis. Z scores (top axis; bars) and P-values (bottom axis; blue line) were computed using permutation testing. The biological process that each condensate is involved in, was manually mapped, and is shown on the left side of the figure. **(I)** Boxplot denoting the number of tissues that a protein is found expressed in. n denotes the number of proteins in each class. P-value computed using Wilcoxon rank sum test.

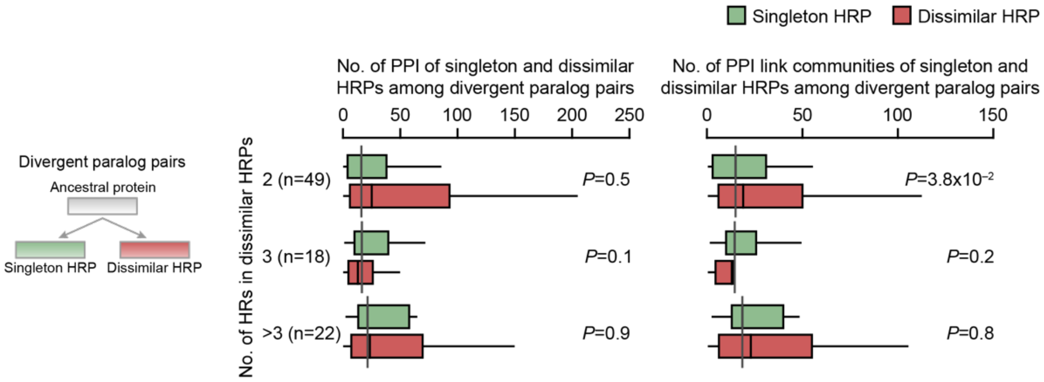


**Supplementary Fig. 2. The change in interactability of dissimilar HRPs is not proportionate with the number of HRs.** Boxplot showing the number of protein interactors (left panel) and the number of link communities (right panel) that singleton and dissimilar HRPs of divergent paralog pairs participate in. P-value computed using Wilcoxon Signed-Rank test. n denotes the number of proteins in each class.

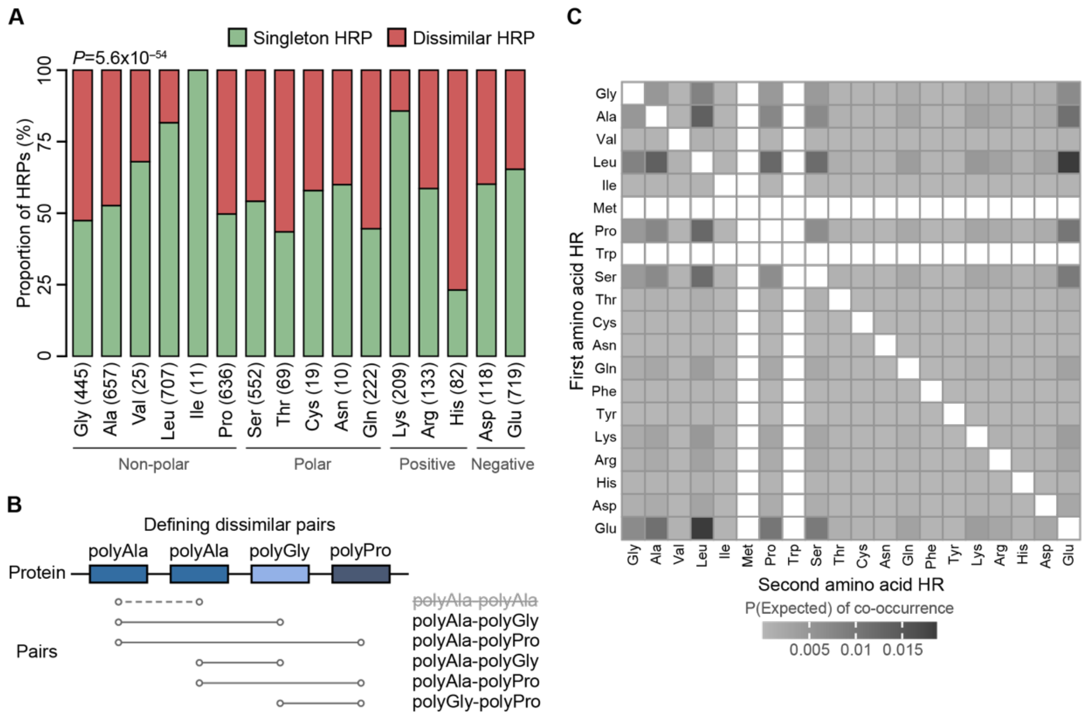


**Supplementary Fig. 3. Non-random distribution of dissimilar HR pairs. (A)** Bar plot showing the proportion of singleton and dissimilar HRPs having different amino acid HRs. n denotes the total number of singleton and dissimilar HRPs of each amino acid HR type. P-value was estimated using Chi-squared test. **(B)** Schema depicting identification of dissimilar HR pairs in a dissimilar HRP with polyAla-polyAla-polyGly-polyPro HRs. **(C)** Heatmap showing the null expectation of co-occurrence of HRs in the human proteome, computed using singleton HRP estimates. P stands for probability.

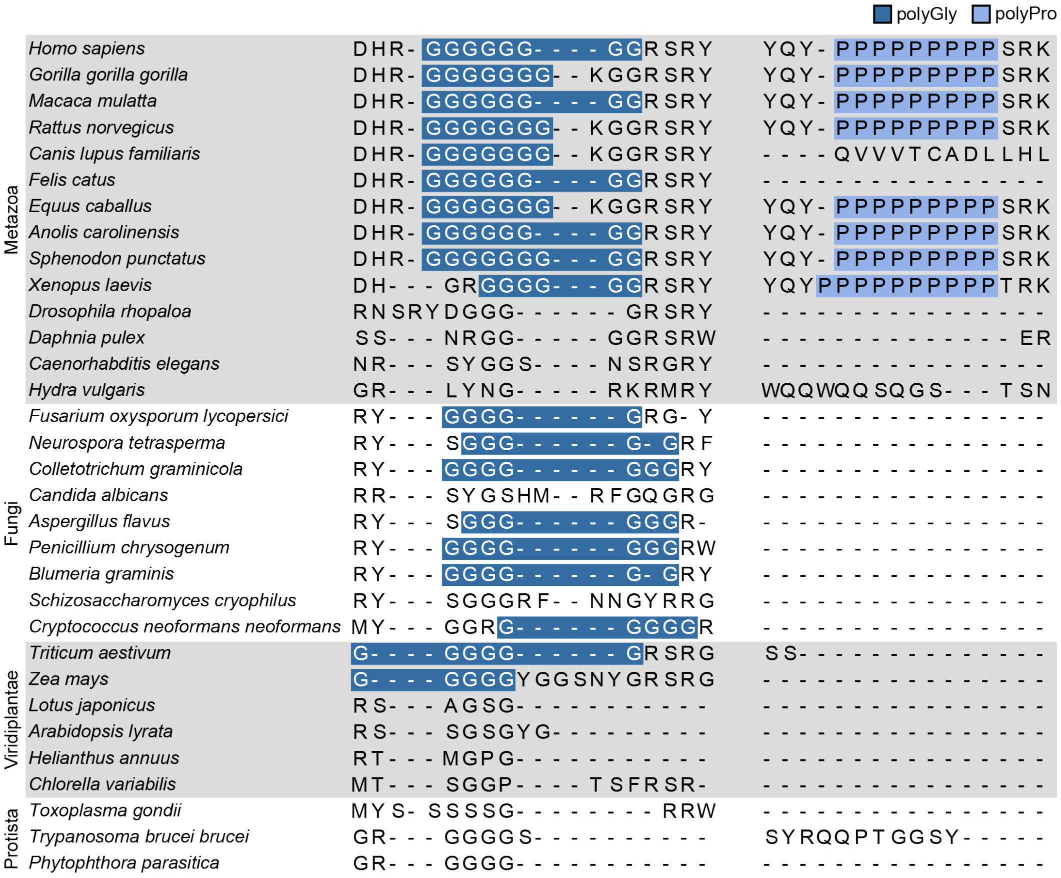

**Supplementary Fig. 4. Conservation profile of the polyGly and polyPro HRs in DDX17.** Multiple sequence alignment of polyGly and polyPro HRs of human DDX17 and its orthologs across species ranging from metazoans to protists (shown on left).

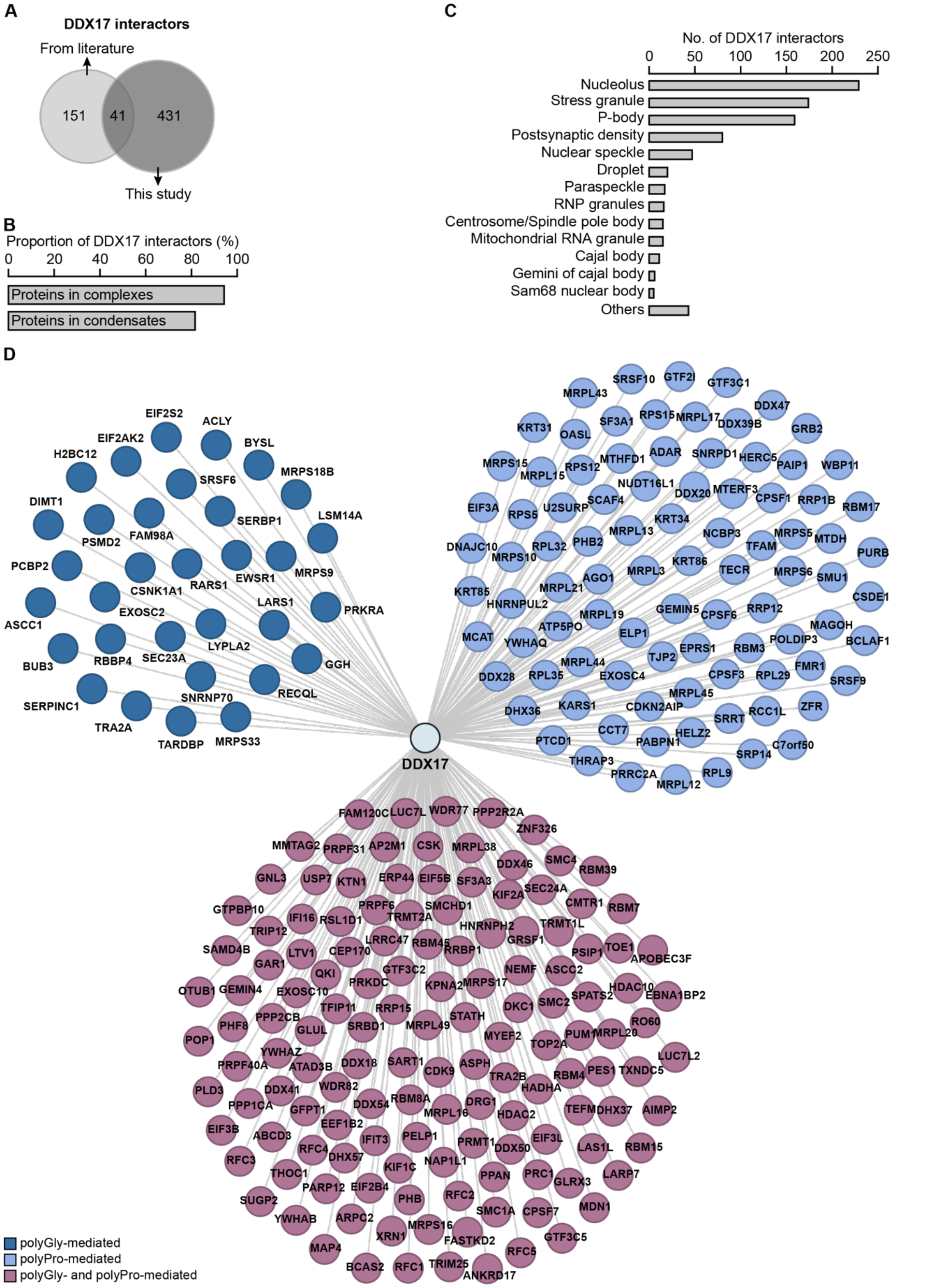


**Supplementary Fig. 5. Interactors of DDX17 and its HRs. (A)** Venn diagram showing the overlap of the protein interactors of DDX17 reported in literature and those identified from our study. **(B)** Proportion of protein interactors of DDX17 that participate in protein complexes and in phase-separated condensates. **(C)** Number of protein interactors of DDX17 that participate in different phase-separated condensates. **(D)** Network showing all protein interactors of polyGly and polyPro HRs of DDX17.

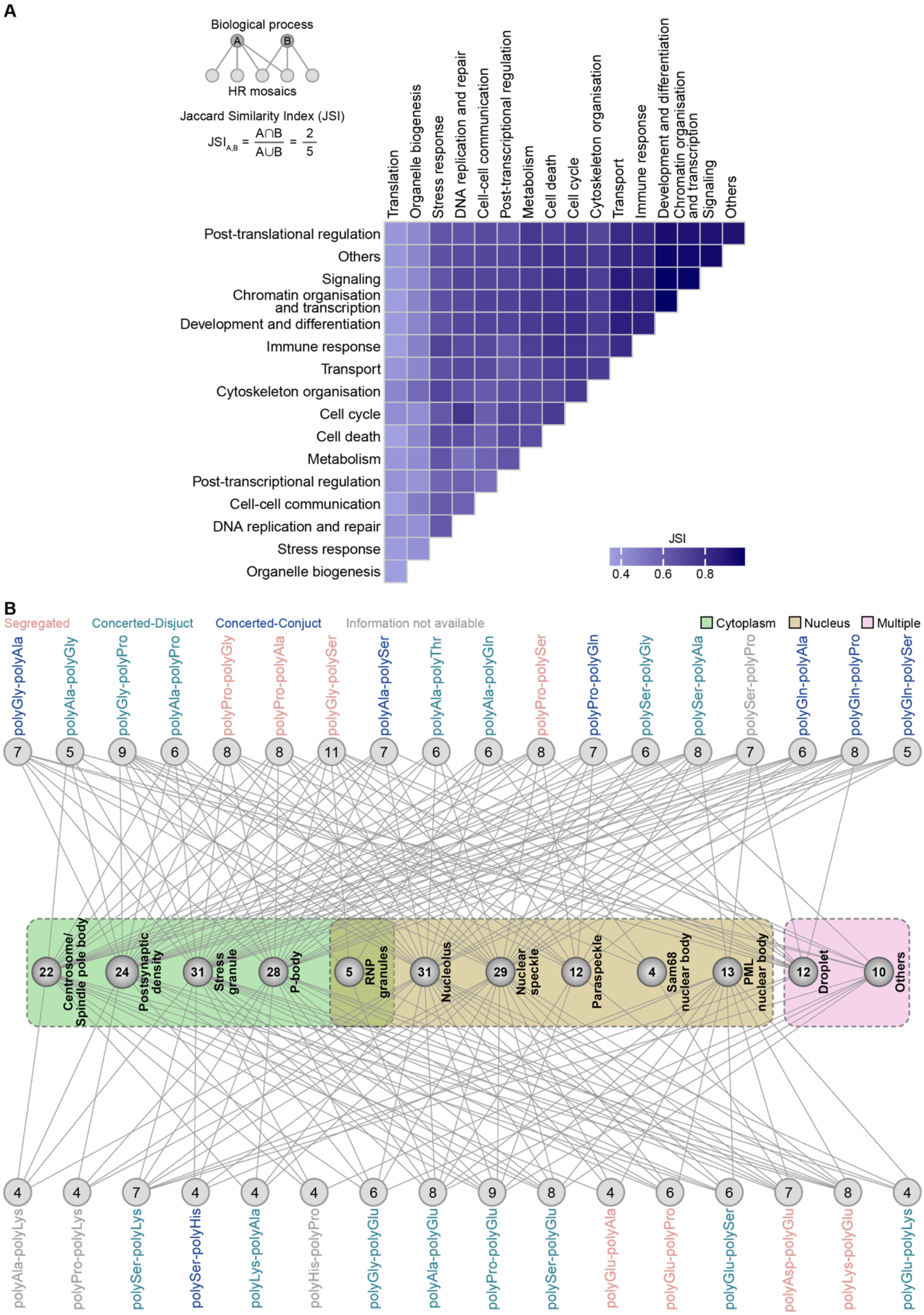


**Supplementary Fig. 6. Shared and specific HR-mosaics across biological processes and condensates. (A)** Heatmap showing Jaccard Similarity Index (JSI) of shared dissimilar HR mosaics between any two biological processes. **(B)** Network representing the different HR mosaics that participate in different phase-separated structures. The number in the HR mosaic nodes denotes the number of phase-separated condensates they participate in. The number in the phase-separated condensate nodes denotes the number of HR mosaics that participate in them.

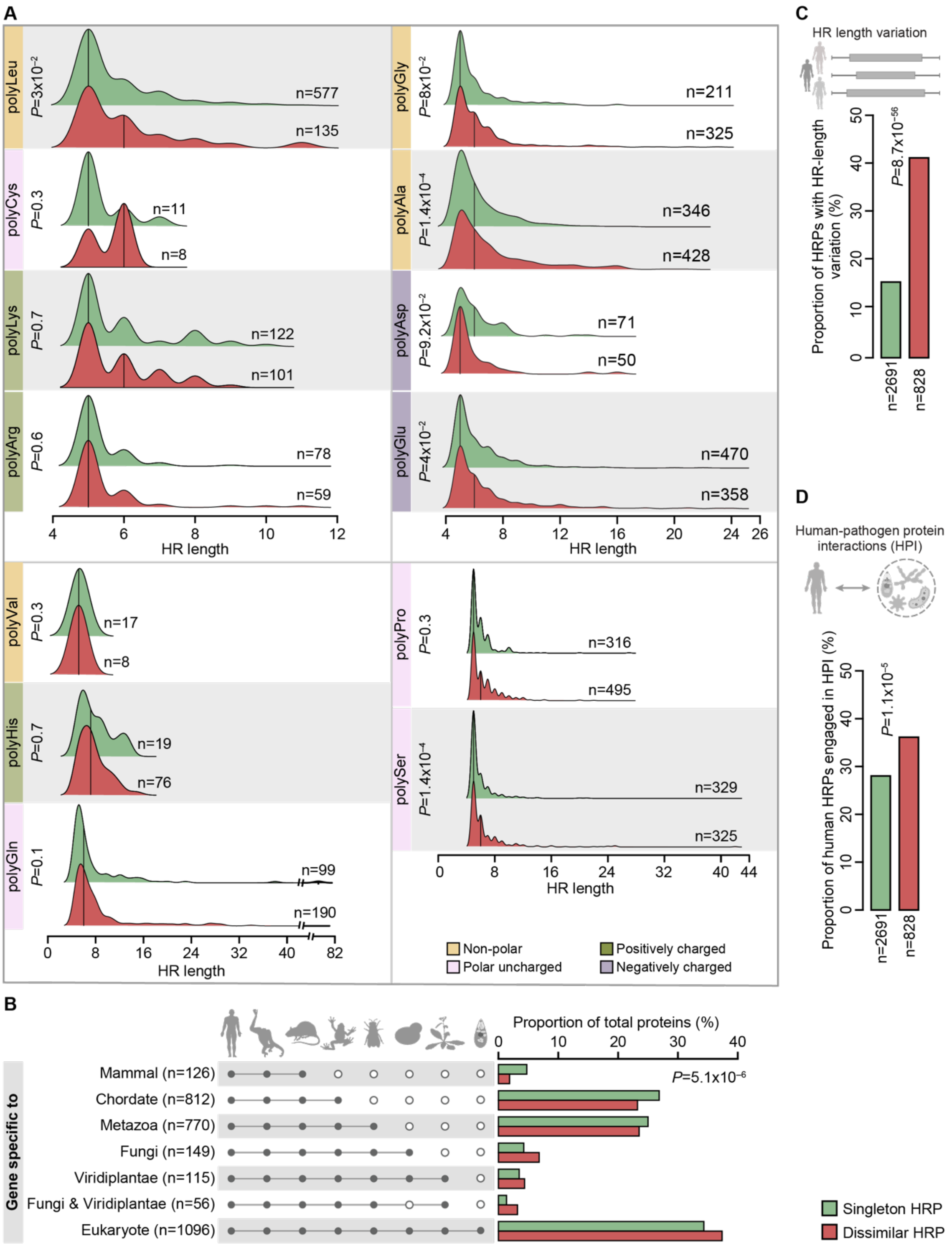


**Supplementary Fig. 7. Distinct attributes of HRs in Dissimilar HRPs. (A)** Ridgeline plot comparing the distribution of the HR lengths by specific type of amino acid HRs between singletons and dissimilar HRPs. The area under the curves represents the density of the distribution of HR lengths. The line within each curve represents the median HR length. n represents the total number of proteins in each category. P-value was computed using Wilcoxon rank-sum test and then corrected for multiple testing using FDR. Amino acid HR types with fewer than five HRs in singleton and dissimilar HRPs were excluded from the analysis. **(B)** Bar plot showing the gene age of human singleton HRPs and dissimilar HRPs. n denotes the number of proteins in each class. P-value was computed using Chi-squared test. **(C)** Proportion of singletons and dissimilar HRPs with length-variable HRs within the human population. n denotes the number of proteins in each class. P-value was computed using Chi-squared test. **(D)** Proportion of singleton and dissimilar HRPs involved in human-pathogen protein interactions (HPI). n denotes the number of proteins in each class. P-value was computed using Chi-squared test.

**Supplementary table S1. List of primers used for generating DDX17 constructs.**

| **Primer** | **Sequence** |
|---|---|
| DDX17WT Forward primer | 5' GAATTCAATGCCCACCGGCTTTGT 3' |
| DDX17WT Reverse primer | 5' GGATCCTTTACGTGAAGGAGGA 3' |
| DDX17ΔG Forward primer | 5' AAGGGTGGTCGTTCTCGT 3' |
| DDX17ΔG Reverse primer | 5' TCTGTGGTCCACAAGCTG 3' |

**Supplementary table S2. List of primers used for generating GFP-HR chimeras.**

| Recombinant construct | Construct label | Forward primer (5'–3') | Reverse primer (5'–3') |
|---|---|---|---|
| pCDNA4-GFP | GFP | GATCGGATCCACCATGGTGAGCAAGGGCGAG | GTACCTCGAGTCACTTGTACAGCTCGTCCATGCC |
| pCDNA4-N-Ala5-GFP | N-$Ala_5$ | GATCGGATCCACCATGGCCGCTGCCGCTGCCGTGAGCAAGGGCGAGGAGCTG | GTACCTCGAGTCACTTGTACAGCTCGTCCATGCC |
| pCDNA4-GFP-C-Ala5 | C-$Ala_5$ | GATCGGATCCACCATGGTGAGCAAGGGCGAG | GTACCTCGAGTCAGGCAGCGGCAGCGGCCTTGTACAGCTCGTCCATGCC |
| pCDNA4-N-His5-GFP | N-$His_5$ | GATCGGATCCACCATGCACCATCACCATCACGTGAGCAAGGGCGAGGAGCTG | GTACCTCGAGTCACTTGTACAGCTCGTCCATGCC |
| pCDNA4-GFP-C-His5 | C-$His_5$ | GATCGGATCCACCATGGTGAGCAAGGGCGAG | GTACCTCGAGTCAGTGATGGTGATGGTGCTTGTACAGCTCGTCCATGCC |
| pCDNA4-N-Ala15-GFP | N-$Ala_{15}$ | GATCGGATCCACCATGGCCGCTGCCGCTGCCGCTGCCGCTGCCGCTGCCGCTGCCGCTGCCGTGAGCAAGGGCGAGGAGCTG | GTACCTCGAGTCACTTGTACAGCTCGTCCATGCC |

| Recombinant construct | Construct label | Forward primer (5'–3') | Reverse primer (5'–3') |
|---|---|---|---|
| pCDNA4-GFP-C-Ala15 | C-$Ala_{15}$ | GATCGGATCCACCATGGTGAGCAAGGGCGAG | GTACCTCGAGTCAGGCAGCGGCAGCGGCAGCGGCAGCGGCAGCGGCAGCGGCAGCGGCCTTGTACAGCTCGTCCATGCC |
| pCDNA4-N-His15-GFP | N-$His_{15}$ | GATCGGATCCACCATGCACCATCACCATCACCATCACCATCACCATCACCATCACCATCACGTGAGCAAGGGCGAGGAGCTG | GTACCTCGAGTCACTTGTACAGCTCGTCCATGCC |
| pCDNA4-GFP-C-His15 | C-$His_{15}$ | GATCGGATCCACCATGGTGAGCAAGGGCGAG | GTACCTCGAGTCAGTGATGGTGATGGTGATGGTGATGGTGATGGTGATGGTGATGGTGCTTGTACAGCTCGTCCATGCC |
| pCDNA4-N-Ala5-GFP-C-His5 | N-$Ala_{5}$–C-$His_{5}$ | GATCGGATCCACCATGGCCGCTGCCGCTGCCGTGAGCAAGGGCGAGGAGCTG | GTACCTCGAGTCAGTGATGGTGATGGTGCTTGTACAGCTCGTCCATGCC |
| pCDNA4-N-Ala15-GFP-C-His15 | N-$Ala_{15}$–C-$His_{15}$ | GATCGGATCCACCATGGCCGCTGCCGCTGCCGCTGCCGCTGCCGCTGCCGCTGCCGCTGCCGTGAGCAAGGGCGAGGAGCTG | GTACCTCGAGTCAGTGATGGTGATGGTGATGGTGATGGTGATGGTGATGGTGATGGTGCTTGTACAGCTCGTCCATGCC |

| **Recombinant construct** | **Construct label** | **Forward primer (5'–3')** | **Reverse primer (5'–3')** |
|---|---|---|---|
| pCDNA4-N-His5-GFP-C-Ala5 | N-$His_5$–C-$Ala_5$ | GATCGGATCCACCAT GCACCATCACCATC ACGTGAGCAAGGGC GAGGAGCTG | GTACCTCGAGTCAG GCAGCGGCAGCGG CCTTGTACAGCTCG TCCATGCC |
| pCDNA4-N-His15-GFP-C-Ala15 | N-$His_{15}$–C-$Ala_{15}$ | GATCGGATCCACCAT GCACCATCACCATC ACCATCACCATCACC ATCACCATCACCATC ACGTGAGCAAGGGC GAGGAGCTG | GTACCTCGAGTCAG GCAGCGGCAGCGG CAGCGGCAGCGGC AGCGGCAGCGGCA GCGGCCTTGTACAG CTCGTCCATGCC |

Other supplementary files associated with this study are listed below.

**Supplementary datasheet 1. All datasets and calculations pertaining to the study. (Provided as an excel sheet)**

**Supplementary movie 1. FRAP time course of DDX17WT protein. (Provided as avi file)**

**Supplementary movie 2. FRAP time course of DDX17ΔG protein. (Provided as avi file)**

**Supplementary movie 3. FRAP time course of DDX17ΔP protein. (Provided as avi file)**

**Supplementary movie 4. FRAP time course of DDX17ΔGΔP protein. (Provided as avi file)**

**Supplementary movie 5. FRAP time course of N-$His_{15}$ chimeric-protein. (Provided as avi file)**

**Supplementary movie 6. FRAP time course of C-$His_{15}$ chimeric-protein. (Provided as avi file)**

**Supplementary movie 7. FRAP time course of N-$Ala_{15}$–C-$His_{15}$ chimeric-protein. (Provided as avi file)**

**Supplementary movie 8. FRAP time course of N-$His_{15}$–C-$Ala_{15}$ chimeric-protein. (Provided as avi file)**
